# Calculating the Effects of Autism Risk Gene Variants on Dysfunction of Biological Processes Identifies Clinically-Useful Information

**DOI:** 10.1101/449819

**Authors:** Olivia J. Veatch, Diego R. Mazzotti, James S. Sutcliffe, Robert T. Schultz, Ted Abel, Birkan Tunc, Susan G. Assouline, Edward S. Brodkin, Jacob J. Michaelson, Thomas Nickl-Jockschat, Zachary E. Warren, Beth A. Malow, Allan I. Pack

## Abstract

Autism spectrum disorders (ASD) are neurodevelopmental conditions that are influenced by genetic factors and encompass a wide-range and severity of symptoms. The details of how genetic variation contributes to variable symptomatology are unclear, creating a major challenge for translating vast amounts of data into clinically-useful information. To determine if variation in ASD risk genes correlates with symptomatology differences among individuals with ASD, thus informing treatment, we developed an approach to calculate the likelihood of genetic dysfunction in Gene Ontology-defined biological processes that have significant overrepresentation of known risk genes. Using whole-exome sequence data from 2,381 individuals with ASD included in the Simons Simplex Collection, we identified likely damaging variants and conducted a clustering analysis to define subgroups based on scores reflecting genetic dysfunction in each process of interest to ASD etiology. Dysfunction in cognition-related genes distinguished a distinct subset of individuals with increased social deficits, lower IQs, and reduced adaptive behaviors when compared to individuals with no evidence of cognition-related gene dysfunction. In particular, a stop-gain variant in the pharmacogene encoding cycloxygenase-2 was associated with having an IQ<70 (i.e. intellectual disability), a key comorbidity in ASD. We expect that screening genes involved in cognition for deleterious variants in ASD cases may be useful for identifying clinically-informative factors that should be prioritized for functional follow-up. This has implications in designing more comprehensive genetic testing panels and may help provide the basis for more informed treatment in ASD.

## Introduction

Autism spectrum disorders (ASD) are a group of neurodevelopmental conditions characterized by core symptoms that include impairments in social interactions, delays in language development and expression of repetitive interests and/or behaviors(*1*). ASDs manifest along a wide distribution of core symptom severity, and numerous different comorbidities are highly prevalent [e.g., intellectual disability(*2*), gastrointestinal issues(*3*)]. Evidence supports contributions from different types of common and rare genetic variation – including inherited and *de novo* single nucleotide variants (SNVs), small insertions or deletions (In/Dels), and large insertions or deletions (CNVs) – in hundreds of genes(*4, 5*). The already large, and rapidly expanding, landscape of genetic factors involved in expression of ASD makes it difficult to determine how results from genetic studies can translate into clinically-useful information(*6-8*). A crucial step toward using genetics to inform more effective, personalized approaches for treatment of individuals with ASD is to better understand how variation in implicated genes influences expression of core symptoms and comorbidities.

While there are more than one hundred implicated genes, many function in the same biological process(*9, 10*). Dysfunction in genetic mechanisms encoding different biological functions may contribute independently to increase risk for ASD. For example, one study observed that a subset of individuals with ASD had *de novo* and rare, inherited variants in synaptic genes but not chromatin modification genes, while another subset had these types of variants in chromatin modification genes but not synaptic genes(*11*). If some individuals with ASD have dysfunction in a particular biological process while others have dysfunction in a separate process, then it may be possible to use genetic data to inform a more personalized (i.e., precision medicine) approach to treatment of symptoms. However, the study mentioned above, and others, have not observed a relationship between genetic and phenotypic differences(*11, 12*). As such, it is difficult to determine if distinguishing dysfunction across different underlying biological processes is clinically useful. Notably, previous studies have focused largely on evaluating contributions from specific *types* of genetic variants (e.g., solely *de novo* and rare variants, or common variants) to explain phenotypic differences(*11-16*). A more holistic approach that incorporates all relevant risk variation is better situated to ask how overall genetic risk is related to particular symptom profiles that are unique to the individual(*17, 18*).

Furthermore, to enable use of disparate genetic information in personalized medicine approaches for ASD, ability to predict functional effects of a given variant on the ASD risk gene and encoded protein is essential and may require functional analysis to test(*19*). While functional study of every suspected ASD risk variant is desirable in the long-term, reliance on such a strategy is not feasible if genetic findings are to be rapidly translated in the clinic. It may be more immediately useful to have computational approaches which incorporate evidence from multiple sources to allow for more thorough *in silico* predictions from patient data to help pinpoint specific genes and variants that should be prioritized for functional follow-up(*20-22*).

To determine if current genetic evidence could help explain variability in ASD symptoms, and ultimately inform treatment approaches, we developed an approach to calculate the likelihood that a biological process with overrepresentation of ASD candidate genes is dysfunctional. We evaluated the approach using whole-exome sequencing and phenotype data from the Simons Simplex Collection (SSC)(*23*). We hypothesized that incorporating evidence from all possible types of genetic variation to calculate cumulative risk of dysfunction overall in biological processes would identify underlying mechanisms contributing to differences in symptomatology among individuals with ASD. We also expected that careful evaluation of the current genetic evidence would be useful to recognizing ASD-related variants that are already clinically-actionable as many individuals carry pharmacogenetics variants which influence how a patient responds to a drug(*24*).

## Materials and Methods

### Identification of Genetic Mechanisms Relevant to ASD

To assess the influences of predicted dysfunction in overall biological processes, we compiled a list of ASD candidate genes using the Autism Informatics Portal (AutDB, http://autism.mindspec.org/autdb/Welcome.do)(*25*), which is continuously updated with manual annotations as new scientific literature is published. As the goal was to determine if any genes evidenced to have a relationship with ASD were useful to understanding symptom variability and informing personalized treatment approaches, all genes were considered regardless of the strength of evidence supporting an association with ASD (December 2017 update). Official Gene Symbols were converted to Ensembl IDs using the Gene ID Conversion Tool available in the database for annotation, visualization and integrated discovery (DAVID)(*26*). Ensembl IDs for a subset of these genes could not be converted via DAVID and were manually identified by searching the Ensembl database. Gene set overrepresentation analyses were run on all candidate genes for ASD using the classic algorithm and Fisher’s exact test from the TopGO package in R(*27*). Overrepresented processes were interrogated to identify terms representing processes useful to ASD etiology (‘unique terms’; *Table S1*). Processes were considered biologically meaningful unique terms if they represented the initial process in each GO hierarchy that was system-, organ-, tissue-, or organelle-specific (e.g., ‘GO:0007399 = nervous system development’). GO term definitions were based on AmiGO version 1.8, GO version 2018-01-01.

### Calculation of Overall Biological Process Dysfunction

Variants identified using whole-exome sequencing available for a total of 2,392 individuals with ASD whose data were included in the Simons Simplex Collection (SSC) dataset(*23*) are provided by the Simons Foundation Autism Research Initiative and WuXi NextCODE: A Contract Genomics Organization (https://www.wuxinextcode.com/). The SSC represents the largest collection of simplex autism families, with one affected child and at least one unaffected sibling, collected to date(*23*). Data are made available to approved researchers via the Sequence Miner Tool 5.24.7. Gender discrepancies were first identified using the ‘Sex Check’ report builder in Sequence Miner. This algorithm evaluates both the ratio of heterozygous SNPS on the X chromosome compared to autosomes and coverage of the Y chromosome gene, *SRY*. Seven individuals with unclear gender assignments, 2 individuals with 47,XYY and one individual with 47,XXX were excluded from analyses. Genome-wide genotyping and whole-exome sequence data for all but one individual in the evaluated dataset (n = 2,381) was previously interrogated to identify *de novo* and rare, inherited copy number variants (CNVs)(*11, 28*). The final analysis dataset included 2,381 individuals who were 4-18 years old at the time of data collection. The dataset was 86% male and 79% parent-reported white (*Table S2*).

Variation Annotation queries were performed in Sequence Miner (*29*) to identify single nucleotide variants (SNVs) and short insertions or deletions (<200bp; In/Dels) located in protein coding gene transcripts that had Variant Effect Predictor(*29*) consequences that were highly likely to be damaging to the encoded protein product (i.e., splice site alterations, gains or losses of stop codons, loss of start codons, or frameshifts). We considered variants were called by either the Genome Analysis Toolkit (GATK)(*30*) or FreeBayes(*31*) software across all 22 autosomes and both sex chromosomes. Quality Control thresholds included depth ≥8 reads and genotype quality for variant calls of ≥20(*32*). Variants flagged as ‘LowQuality’ as indicated by GT Filter criteria were excluded.

The final list of variants passing QC, that were predicted by VEP to be very likely to be damaging, were interrogated to identify those located in transcripts for ASD risk genes (included the Autism Informatics Portal) that were protein coding (*Table S3*). Notably, the Sequence Miner platform reports Ensembl IDs for each gene in the query output. While these were used to ensure the appropriate VEP predictions and help search the current Ensembl database, some of the Ensembl IDs provided with this platform were outdated and are now represented by new IDs. As such, both Ensembl IDs and gene names were cross-referenced to compare those provided by Sequence Miner and those mapped using DAVID and manual searches. Discrepancies were further interrogated to verify that the VEP prediction was not based on a variant location in an alternate transcript that is not supported by evidence in Ensembl.

There is substantial variability in pathogenicity predictions depending on the algorithm employed (e.g., based on variant location, evolutionary conservation, protein structure/function)(*21, 33*). Therefore, to more completely assess the likelihood of a variant being damaging and ultimately resulting in a dysfunctional protein product, nine different variant prediction algorithms were run on all of the variants pulled from Sequence Miner using filter-based annotation from ANNOtate VARiation (ANNOVAR) software(*34*). *In silico* prediction algorithms included: 1) Sorts Intolerant From Tolerant (SIFT)(*35*), 2) Polymorphism Phenotyping v2 (Polyphen-2) HVAR(*36*), 3) Mutation Taster(*37*), 4) Mutation Assessor(*38*), 5) Likelihood Ratio Test (LRT)(*39*), 6) FATHMM-MKL(*40*), 7) PROVEAN, 8) MetaLR(*41*), and 9) Mendelian Clinically Applicable Pathogenicity (M-CAP)(*42*). Genomic locations of variants available from Sequence Miner are based on Human Genome Build GRCh37/hg19; all analyses were conducted based on these genomic locations. Each prediction algorithm uses different nomenclature to denote variant predictions. To allow for cross-comparison of results from different predictors, scores were recoded as either benign (B), damaging (D), or unknown (U) as follows: SIFT: damaging (D) = D, tolerant (T) = B; Polyphen2 HVAR probably damaging (D) = D, possibly damaging (P) = U, benign (B) = B; LRT: deleterious(D) = D, unknown(U) = U, neutral(N) = B, Mutation Taster: disease causing automatic(A) = D, disease causing(D) = D, polymorphism(N) = B, polymorphism automatic(P) = B; Mutation Assessor: predicted functional (H, M) = D, predicted non-functional (L, N) = B; FATHMM-MKL: damaging(D) = D, neutral(N) = B; PROVEAN: deleterious(D) = D, neutral(N) = B; MetaLR: damaging(D) = D, tolerant(T) = B; M-CAP: pathogenic(D) = D, benign(T) = B.

We developed the following equation to calculate the likelihood that a variant was damaging to the function of the encoded protein product:

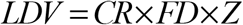

Where ***LDV*** = the likelihood that the variant is damaging; ***CR*** = the number of variant callers that called the variant (based on GATK and FreeBayes software); ***FD*** = ((***D-B***) + 1))/(***N*** + 1) where ***D*** = the number of *in silico* prediction algorithms that called the variant damaging, ***B*** = the number of algorithms that called the variant benign, ***N*** = the total number of algorithms that provided a prediction for the variant, and 1 = a constant to account for the fact that variants were preselected according to variant effect predictions indicating a high potential to be deleterious to at least one gene transcript based on the genetic location; and ***Z*** = zygosity where heterozygous calls = 1 and homozygous calls = 2. To reduce the likelihood of false positive calls overly influencing genetic risk scores, variants were weighted such that if only one variant caller recognized the base pair alteration compared to reference ***CR*** = 0.5. If the variant was called by both the GATK and FreeBayes callers ***CR*** = 1. ***FD*** scores ranged from −0.8-1.0; however, as the goal was to identify variants that were more likely to be deleterious, all negative scores were recoded to zero. Regarding zygosity for sex chromosomes, as it is difficult to determine which X chromosome is inactivated using the data available, female individuals with heterozygous variants on the X chromosome were weighted the same as autosomal variants. In addition, for X chromosome variants called as heterozygous in males, those located within Pseudoautosomal Regions (PAR) were weighted the same as autosomal variants. Male heterozygous X chromosome variants located outside of PAR1 and PAR2 were considered homozygous and were weighted as such in genetic risk scores.

Hg19 genomic locations of rare, inherited and validated, *de novo* CNVs previously reported in Sanders et al., 2015(*11*) and Krumm et al., 2015(*28*) that encompassed coding and regulatory regions of protein coding transcripts for ASD candidate genes were pulled from supplemental data included in these publications. Bedtools(*43*) was used to identify regions of overlap between CNVs reported across the previously published studies. Gene-based annotations in ANNOVAR were used to identify CNVs that encompassed portions of the coding (i.e., exonic, splice-site) and proximal promoter (i.e., 5’-UTR) regions (*Tables S4-S5*). CNVs were given weights equal to SNVs and In/Dels with the strongest likelihood of being damaging based on the distribution of ***FD*** scores described above and variant weights for all CNVs were set equal to 1. The currently published data for CNVs report only the presence of a deletion or duplication in a particular genomic region but not the predicted number of copies; however deletions were expected to occur on only one chromosome(*11, 28*). Deletions and amplifications were assumed to occur on only one chromosome. In addition, while some CNVs were not reported for the same evaluated individual, the analysis datasets across the two prior publications did not completely overlap. Therefore, whether or not both studies reported the CNV was not included in variant weights.

Separate genetic risk scores were then calculated for each individual to assess the likelihood of dysfunction in overall genetic mechanisms that represented unique GO-defined biological processes with overrepresentation of ASD candidate genes. We developed the following equation to calculate the likelihood of genetic dysfunction in biological processes:

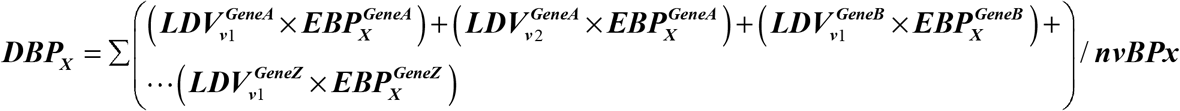

Where ***DBP_X_*** = Dysfunction of Biological Process X and is the sum of the products of 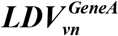 = the likelihood that variant ***n*** is damaging to gene A, and 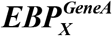 = the sum of the frequencies of the GO evidence codes, across all genes assigned to biological process X, that were used to assign gene A to biological process X (*Fig. S1*) plus the number of assigned child terms for biological process X, divided by the total number of child terms available for biological process X. **nvBPx** = the number of variants assigned to biological process X. We expected that a gene having more than one likely damaging variant was increased evidence that the encoded protein product was dysfunctional. Furthermore, the size of the transcripts was not correlated with the number of variants identified in the gene (R^2^ = 2.0×10^−4^). Therefore, we did not correct for multiple variants per gene.

### Clustering of Biological Process Dysfunction Scores

To cluster individuals based on overall genetic risk, we used an approach that we previously developed and showed was capable of identifying genetically-meaningful subgroups in ASD (*44*). Briefly, the correlation structure across the genetic risk scores was determined by calculating pairwise Spearman’s rank correlation coefficients. As score ranges varied by biological process, all scores were transformed into Hazen percentile ranks to be more comparable. To help ensure that correlated genetic risk scores did not overly influence results, Gower dissimilarity matrices were calculated using correlation-based weights with the ‘FD’ package v1.0-12 in *R(45).* The threshold for non-independence of genetic risk scores was p□≥□0.50, or moderate to strong correlation(*46*). Correlated scores were weighted to allow for only partial contributions to analyses. The ‘clValid’ package v0.6-6 in R was used to evaluate different methods for internal validity using connectivity, silhouette width, and the Dunn index while partitioning the dissimilarity matrix into anywhere from 2 to 5 clusters(*47*). Clustering methods that are available for evaluation in the clValid package include: 1) agglomerative hierarchical, 2) partitioning around medoids, 3) self-organizing tree algorithm, 4) model-based, 5) divisive hierarchical, and 6) fuzzy k-means. The final clustering solution was performed using the agglomerative hierarchical method via the ‘cluster’ package v2.0.7-1 in *R*(*48*). Final cluster solution validity was assessed by performing 1,000 data permutations and comparing clustering of real versus permuted genetic risk scores with the Adjusted Hubert-Arabie Rand *index(49*). Sensitivity and regression analyses were performed to determine if dysfunction in any particular biological process was important to definition of the final cluster solutions. Chi-square tests were used to determine if having variants with ***LDV>*** 0 in any particular gene was associated with assignment of individuals to genetic clusters.

### Differences in ASD-Related Phenotype Variables Based on Genetic Subgroup

Phenotype variables representing quantitative or ordinal severity measures for symptoms assessed in the SSC standard phenotype battery and medical history intakes were downloaded directly from SFARI Base (https://base.sfari.org/) and were available for the majority of the ASD probands included in the genetic data analyses (99.66%, n = 2,373). For more information on symptom severity measurements used for the SSC see Fischbach and Lord, *2010(23).* When available, normalized z-scores or age-standardized scores were used. Head circumferences were transformed to *z*-scores by standardizing for age and sex using a typically developing population(*50*). Sleep duration was determined using current answers to the question “On average, how many hours/night [does your child sleep]?” obtained from the medical history intakes as described in our previous study(*51*). Student’s t-tests were used to compare mean scores for symptom severity measures, that were available for at least half of the analysis dataset, between the individuals assigned to genetic clusters. Age was not associated with measures that were significantly different between clusters (p≥0.43). For measures with sex-specific differences, additional t-tests were conducted that were stratified by sex. Chi-square tests were used to determine if having variants with ***LDV>*** 0 in any particular gene was associated with assignment to the genetic clusters. Logistic regression was used to test if having variants with ***LDV>0*** in cluster-associated genes was associated with: 1) individuals with ASD compared to unaffected siblings in all races and only in white individuals, 2) increased risk for intellectual disability (IQ<70) or reports of irritable bowel syndrome while adjusting for gender and race. False discovery rate was controlled for using the Benjamini-Hochberg procedure(*52*).

Principal Component Analysis (PCA) was conducted while applying correlation-based weights to allow only partial contributions of moderately-strongly correlated phenotype variables *(p□≥□0.50),* similar to that described for clustering of genetic risk scores. Phenotype variables were transformed to Hazen percentile ranks prior to PCA. PCA was conducted without scaling as variables did not contribute equal weights. The number of dimensions of the PCA was estimated via cross-validation. PCA was then performed on percentile ranked phenotype data using the ‘FactoMineR’ package v1.41 in R(*53*).

## Results

### Novel Approach Calculates Dysfunction in Biological Processes Underlying ASD

At the time of these analyses, there were 989 different protein coding ASD risk genes included in the Autism Informatics Portal (December 2017 update). 2,482 Gene Ontology (GO)(*54, 55*) biological processes defined for humans were overrepresented for ASD risk genes based on a significance threshold of p<0.05; 16 terms had the lowest possible p-values (p<1×10^−30^; *Fig. 1, Table S1*). Of the 16 top overrepresented terms, four GO terms – nervous system development (GO:0007399), synaptic signaling (GO:0099536), cognition (GO:0050890), and regulation of membrane potential (GO:0042391) – represented unique processes. There were 400 ASD candidate genes with evidence for involvement in at least one of these four biological processes. The genes that remained unassigned to any process were overrepresented in the chromosome organization process (GO:0051276, p = 7.10×10^−12^; *Fig. 1*, *Table S1*). An additional 82 genes were evidenced to be involved in chromosome organization. There were no unique biological processes with evidence of overrepresentation for the remaining 507 unassigned ASD candidate genes (*Table S1, Fig. S2*). The overlap in ASD risk genes assigned the five overrepresented biological processes representing unique terms is shown in Figure S3A.

**Figure 1.**
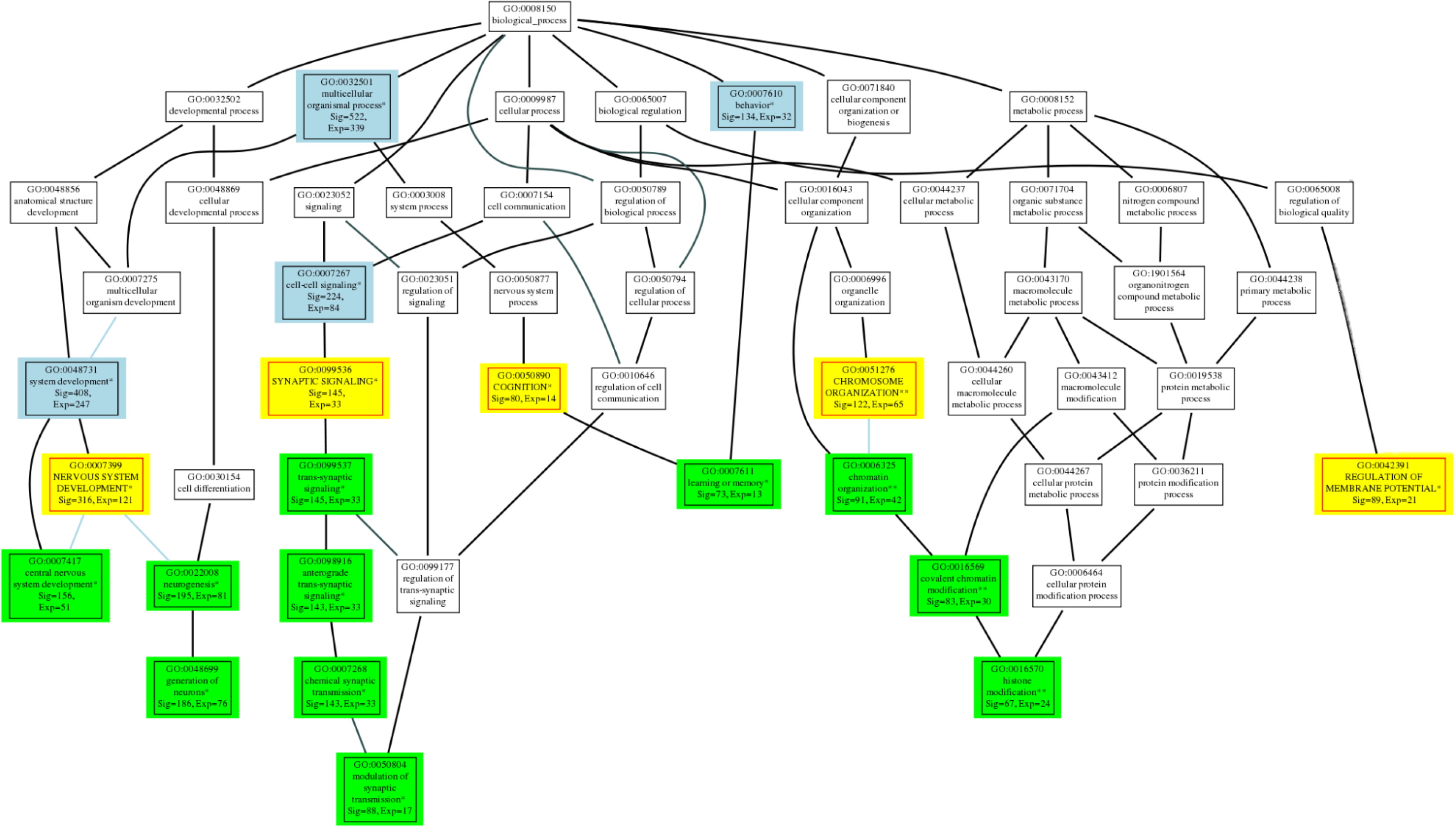
Selection of unique biological processes with overrepresentation of ASD candidate genes for further study. Shown is the distribution of significant terms in the GO structure for biological processes (GO:0008150). Terms highlighted in yellow indicate unique terms selected due to their place in the hierarchy and meaningfulness to ASD etiology. Terms highlighted in blue indicate significant processes considered too broad to be meaningful and green indicates significant child terms with complete genetic overlap to unique terms. Sig = the number of ASD candidate genes assigned to the process, Exp = the expected number of genes assigned by chance. *denotes terms that were significant at Fisher’s exact FDR<1.0×10^−30^ following the primary analysis of all 989 ASD risk genes, ** denotes terms that were significant at Fisher’s exact FDR ranging from 3.5×10^−17^ to 7.1×10^−12^ following the secondary analysis run on genes unassigned to the top processes. Black lines connect terms that regulate each other, blue lines connect terms that are part of each other.

There were 2,077 unique SNVs and In/Dels predicted by Variant Effect Predictor (VEP) to be damaging (*Table S3*). Predictions of variant effects based on nine other algorithms that use information in addition to genetic location indicated that 730 of the 2,077 variants were more often predicted damaging compared to benign (i.e., ***LDV*** >0). The majority of the individuals in the analysis dataset (n = 2,295, 96.35%) had a variant with ***LDV*** >0 in an ASD risk gene. On average, there were ∼15 variants [µ = 14.6(5.3)] observed per individual that was predicted to be damaging more often than benign, and ∼11 different [µ = 11.3, (4.2)] ASD candidate genes per person with possibly damaging variants. None of the variants that were *de novo* were predicted to be benign and inherited variants were more often predicted to be damaging if the consequence related to a frameshift, splice-site alteration, or incorporation of a premature stop codon (*Fig. 2*). Screening data reported in previous studies(*11, 28*) for *de novo* and rare, inherited structural variation in the SSC dataset identified 572 unique Copy Number Variants (CNVs) encompassing coding regions or proximal promoter elements of 354 ASD candidate genes (*Tables S4-S5*). There were 546 individuals in the analysis dataset with ≥1 CNV that was likely to cause dysfunction in ≥1 ASD candidate gene; 292 CNVs encompassed more than one gene. In total, there were 751 currently implicated genes with either a SNV, In/Del or CNV with ***LDV*** >0. Of these, 355 were assigned to at least one unique process that was overrepresented for ASD risk genes (*Fig. S3B*).

**Figure 2.**
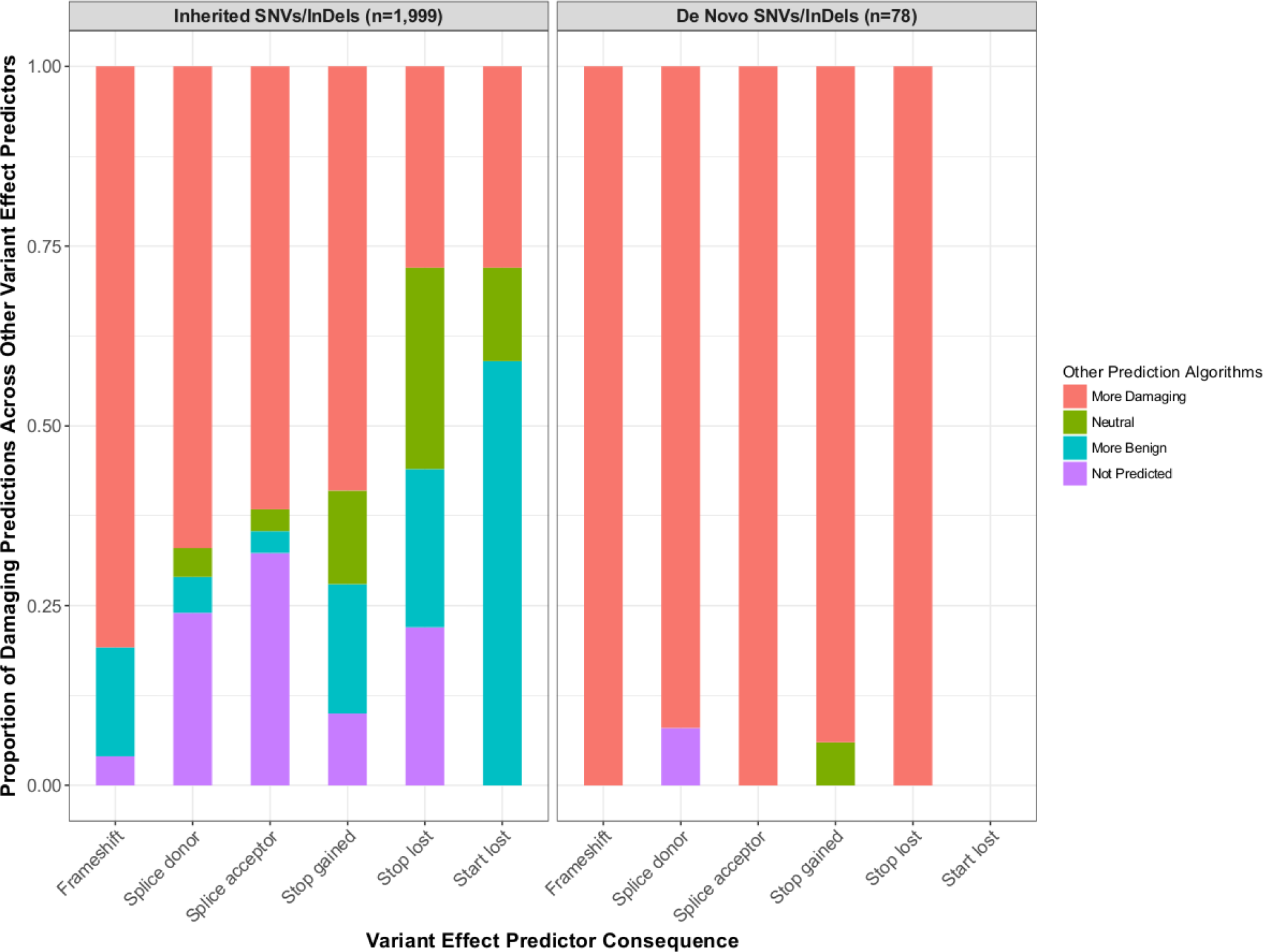
Proportion of VEP consequences predicted to be damaging based on nine prediction algorithms. Inherited variation resulting in frameshifts, splice-site and stop gains were more often predicted damaging compared to benign, while variants predicted to cause the loss of either stop or start sites were equally or more often predicted to be benign. *De novo* variants, regardless of the consequence were more often damaging.

Most individuals in the dataset (98.1%) had evidence indicating genetic dysfunction in more than one of the evaluated biological processes. There were five individuals with evidence for dysfunction only in nervous system development, 35 with evidence for dysfunction only in chromosome organization, and five with no evidence for dysfunction in any of the evaluated processes. Scores for dysfunction in nervous system development, synaptic signaling, and regulation of membrane potential were moderately to strongly correlated. Scores reflecting dysfunction in cognition and chromosome organization were weakly correlated with each other and other scores (*Fig. 3A*).

**Figure 3.**
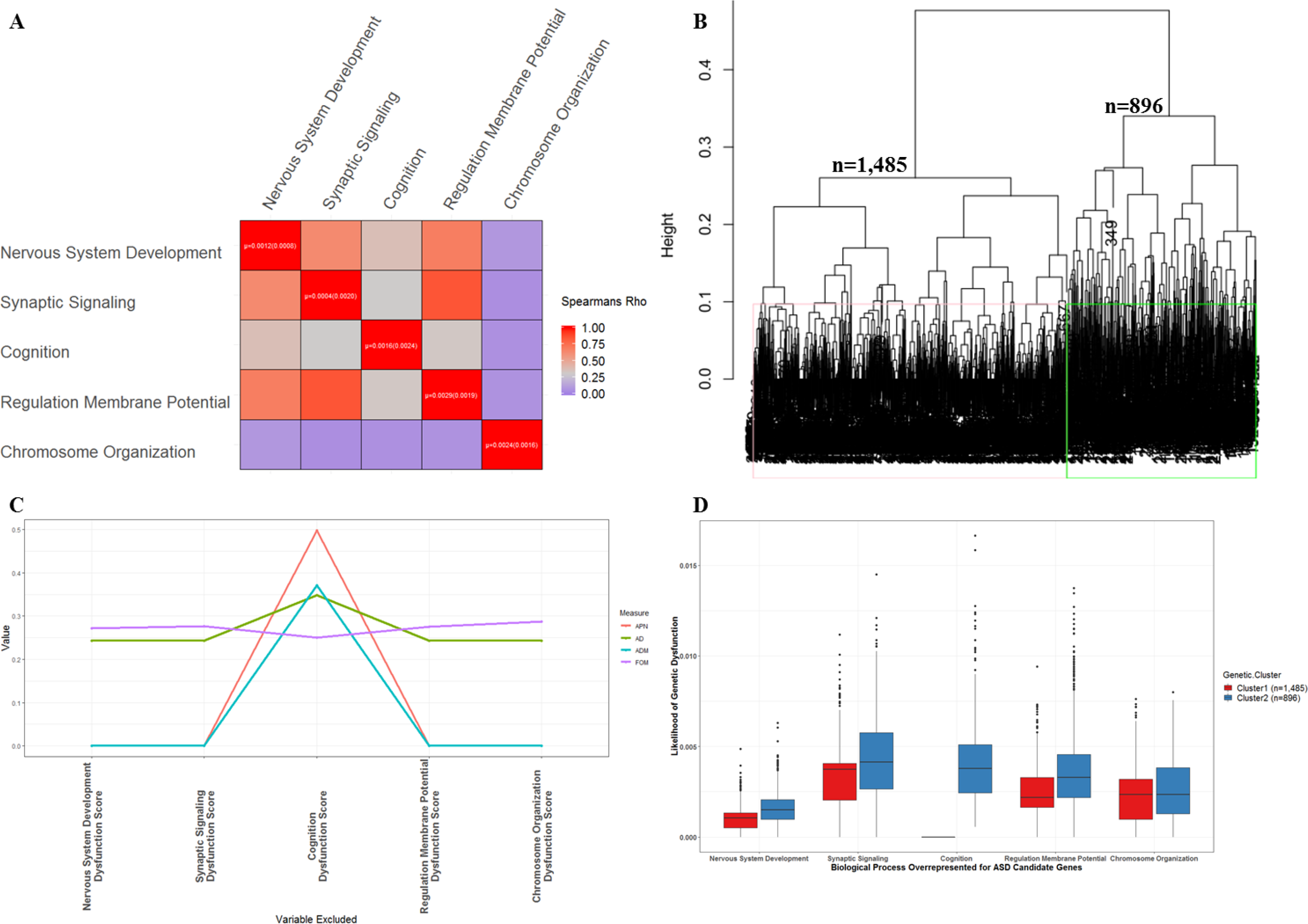
Clustering individuals based on overall biological process dysfunction. **A)** Correlation across scores reflecting dysfunction in biological processes with overrepresentation of ASD candidate genes indicates many individuals have dysfunction in >1 process. **B)** Clustering identified two distinct subgroups of individuals with more similar scores for overall biological process dysfunction (agglomerative coefficient = 0.96). **C)** Sensitivity analyses indicate removing the scores had the strongest effect on stability of the clustering solution. APN = average proportion of non-overlap, AD = average distance, ADM = average distance between means, FOM = figure of merit. **D)** Evidence of dysfunction in genes involved in cognition primarily defined separation of individuals into either cluster 1 (no cognition gene dysfunction) or cluster 2 (cognition gene dysfunction).

A clustering analysis was then performed on ***DBP_X_*** scores reflecting the likelihood of genetic dysfunction in each of the five unique biological processes. Agglomerative hierarchical clustering identified two valid subgroups of individuals (n_Cluster 1_ = 1,485, n_Cluster2_ = 896) (*Fig. 3B, Fig. S4*). This solution was significantly different from clustering permuted datasets (HubertArabieRandIndex = −1.2×10^−4^), further evidence supporting validity of the clustering analysis. Sensitivity analyses indicated that scores reflecting genetic dysfunction in the cognition biological process had the strongest influence on the stability of the clusters (*Fig. 3C*). Notably, all of the individuals assigned to the smaller cluster had evidence of dysfunction in genes involved in cognition (‘cognition gene dysfunction cluster’) while none of the individuals assigned to the larger cluster had evidence for dysfunction in these genes (*Fig. 3D*).

### Three Cognition Genes are Associated with Distinct ASD Genetic Subgroup

Of the 61 cognition genes with likely damaging variants identified in the SSC dataset, there were three genes (*PTGS2*, *ABCA7*, and *SHANK3*) that were strongly associated with assignment to the cognition gene dysfunction cluster (*Table 1A, Table S6*). There were 196 individuals who were heterozygous for a stop-gain variant in exon 4 (rs200314986; transcript ENST00000367468.9:c.366C>A, ENSP00000356438.5:p.Tyr122Ter) of the *prostaglandin-endoperoxide synthase 2* (*PTGS2*) gene, which results in a shortened transcript that is missing the final 6 exons. This variant was more frequent in individuals with ASD compared to unaffected siblings (*Table 1B*). There were 17 different likely damaging variants observed in 280 individuals in the *ATP Binding Cassette Subfamily A Member 7* (*ABCA7*) gene. These included six frameshifts, four splice-sites, four stop-gains, one stop-loss, one inherited deletion of the first 11 exons (CNV size = 18.7kb), and one inherited amplification encompassing exons 27-40 (CNV size = 4.5kb). For the *SH3 and multiple ankyrin repeat domains 3* (*SHANK3*) gene, there were 294 individuals who were heterozygous for a splice-site variant (rs150909992) that changes the 5’ end of an intron in transcript variant ENST00000445220.2, and two individuals with *de novo* CNVs that deleted the entire coding region (CNV sizes>3Mb).

**Table 1A.** Genes associated with the cognition gene dysfunction cluster.

| Genetic Cluster | *PTGS2* |  | *ABCA7* |  | *SHANK3* |  |
| --- | --- | --- | --- | --- | --- | --- |
|  | No Variant | Variant | No Variants | Variants | No Variants | Variants |
| Cluster 1 |  |  |  |  |  |  |
| Observed | 1,485 | 0 | 1,485 | 0 | 1,485 | 0 |
| Expected | 1,363 | 122 | 1310 | 175 | 1,300 | 185 |
| Cluster 2 |  |  |  |  |  |  |
| Observed | 700 | 196 | 616 | 280 | 600 | 296 |
| Expected | 822 | 74 | 791 | 105 | 785 | 111 |
| Total | 2185 | 196 | 2,101 | 280 | 2085 | 296 |
| Pearson $\chi^2$ | 353.98 | | 525.91 | | 560.23 | |
| Fisher's exact | <1.0x10<sup>-32</sup> |  | <1.0x10<sup>-32</sup> |  | <1.0x10<sup>-32</sup> |  |

**Table 1B.** Association of cluster-associated cognition genes with Autism Spectrum Disorder.

| ASD Diagnosis | *PTGS2* |  | *ABCA7* |  | *SHANK3* |  | Total |
| --- | --- | --- | --- | --- | --- | --- | --- |
|  | No Variant | Variant | No Variants | Variants | No Variants | Variants |  |
| All Reported Races |  |  |  |  |  |  |  |
| ASD | 2,185 | 196 | 2,101 | 280 | 2,085 | 296 | 2,381 |
| Unaffected Siblings | 1,700 | 100 | 1,602 | 198 | 1,431 | 396 | 1,800 |
| Odds Ratio (95%C.I.)† | 1.52 (1.18, 1.97) |  | 1.08 (0.88, 1.31) |  | 0.55 (0.46, 0.65) |  |  |
| p-value | 5.0x10<sup>-4</sup> |  | 2.4x10<sup>-1</sup> |  | <1.0x10<sup>-5</sup> |  |  |
| FDR | 7.5x10<sup>-4</sup> |  | 2.4x10<sup>-1</sup> |  | 3.0x10<sup>-5</sup> |  |  |
| Reported White Race |  |  |  |  |  |  |  |
| ASD | 1,644 | 184 | 1,634 | 194 | 1,594 | 234 | 1,828 |
| Unaffected Siblings | 1,280 | 85 | 1,240 | 125 | 1,095 | 270 | 1,365 |
| Odds Ratio (95%C.I.)† | 1.69 (1.28, 2.23) |  | 1.18 (0.92, 1.50) |  | 0.60 (0.49, 0.72) |  |  |
| p-value | 1.0x10<sup>-4</sup> |  | 9.7x10<sup>-2</sup> |  | <1.0x10<sup>-5</sup> |  |  |
| FDR | 1.5x10<sup>-4</sup> |  | 9.7x10<sup>-2</sup> |  | 3.0x10<sup>-5</sup> |  |  |

Of the 61 genes used to calculate *DBP<sub>Cognition</sub>* scores, **A)** three genes were significantly associated with assignment of individuals to the cluster with evidence for dysfunction in cognition genes (i.e., Genetic Cluster 2). **B)** In particular, a variant in *PTGS2* was more frequent in individuals with ASD compared to unaffected siblings. Tests were conducted in all individuals and only in individuals with white race reported to account for potential influences of population stratification. †Odds ratios denote the likelihood for an individual to have an ASD diagnosis given the presence of any likely damaging variant in *SHANK3*, or *ABCA7*, or the T-allele (i.e. a stop-gain variant) in *PTGS2*.

### Individuals with Cognition Gene Variants Have More Severe Symptoms

Among the 27 ASD-related symptom measures that were available for at least half of the dataset (*Table S7*), the severity of social impairment based on teacher reports on Social Responsiveness Scales (SRS-TR), intelligence quotient (IQ) scores, personal and social skills measured using composite standard scores from the Vineland Adaptive Behavior Scales, receptive vocabulary measured via the Peabody Picture Vocabulary Test, and the severity of ASD-related abnormalities exhibited by 36 months of age (i.e., Developmental Abnormality scores) from the Autism Diagnostic Interview-Revised (ADI-R) were different between the genetic clusters (*Fig. 4A, Table S8A*). After false discovery rate corrections, the observations that individuals with dysfunction in cognition genes had increased severity of social impairment reported by teachers on the SRS-TR and reduced IQs remained significant (*Fig 4A, Table S8A*). Notably, both nonverbal and verbal IQ scores were lower in the genetic subgroup defined by dysfunction in cognition genes (*Fig 4A, Table S8B*). Sex-stratified mean comparisons indicated that differences between the genetic clusters for SRS-TR scores and verbal IQs were more significant in males compared to females (*Table S8C*).

**Figure 4.**
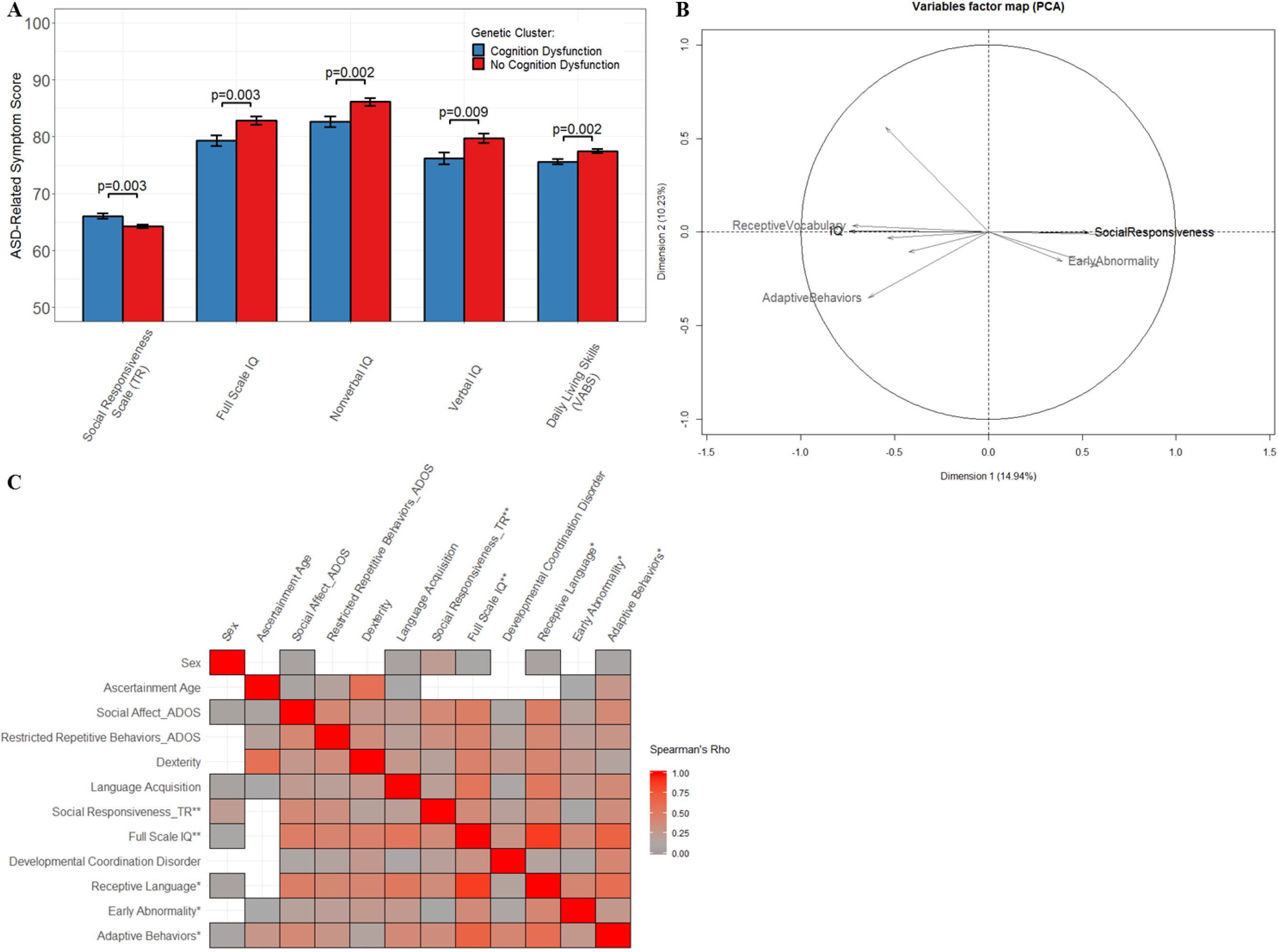
Relationship between genetic and phenotypic differences. **A)** T-tests comparing differences in the 27 quantitative ASD-related symptom measures between genetic clusters identified that social impairment was more severe and IQs and daily living skills were reduced in the cognition gene dysfunction cluster. **B)** Principal components analysis, while adjusting for correlations, of all 27 symptom measures identified that symptoms that were different between the genetic clusters majorly contributed to overall phenotype variability (as defined by Dimension 1). Black indicates symptom differences that remained significant (FDR≤0.04) following multiple testing correction, gray indicates symptom differences based on an unadjusted significance threshold (p≤0.03), and unlabeled arrows indicate symptoms that were not different but had strong contributions to phenotype variability. **C)** Significant (p<0.05) correlations are shown indicating that absolute values for many symptoms that were different between genetic clusters were correlated with those contributing majorly to overall phenotype variability.

To determine how much of the overall variability in ASD symptomatology was explained by symptoms that were different between the genetically distinct subgroups, principal components analysis (PCA) was conducted, while adjusting for correlated variables, on phenotype data from the subset of the dataset with all evaluated measures (n = 543). Five principal components (PCs) were able to define the majority of the variability in symptoms (cumulative percentage of variance = 46.97%). Of the 27 measures evaluated, teacher reports on SRS-TR were the 5^th^ strongest contributor to the cumulative variability defined by the first component of the data (*Fig. 4B, Fig. S5*). The strongest correlation for SRS-TRs (ρ = 0.39) was with the variable which contributed the most to explaining the phenotypic heterogeneity defined by PC1, social and communication impairment observed via the Autism Diagnostic Observation Schedule (*Fig. 4C, Figs. S5-S6*). Full scale IQs were the 6^th^ strongest contributor to the variability defined by PC1 (*Fig. 4B, Fig. S5*). Full scale IQ scores were moderately correlated with scores for dexterity (Purdue Pegboard Test, ρ = 0.45; *Fig 4C*) and language acquisition (non-word repetition task, ρ = 0.48; *Fig. 4C*) which were the 3^rd^ and 4^th^ largest contributors to PC1, respectively (*Fig. S5*). Of the top three genes associated with assignment to the ‘cognition dysfunction cluster’, the stop-gain variant in the *PTGS2* gene was associated with increased risk for having an IQ score reflecting intellectual disability (*Table 2A*) and reports of comorbid irritable bowel syndrome, when adjusting for sex and race (*Table 2B*). The majority of the individuals with ASD, and all of the unaffected siblings with the variant inherited it from at least one parent. There were six individuals with ASD whose parents did not appear to have the variant.

**Table 2A.**
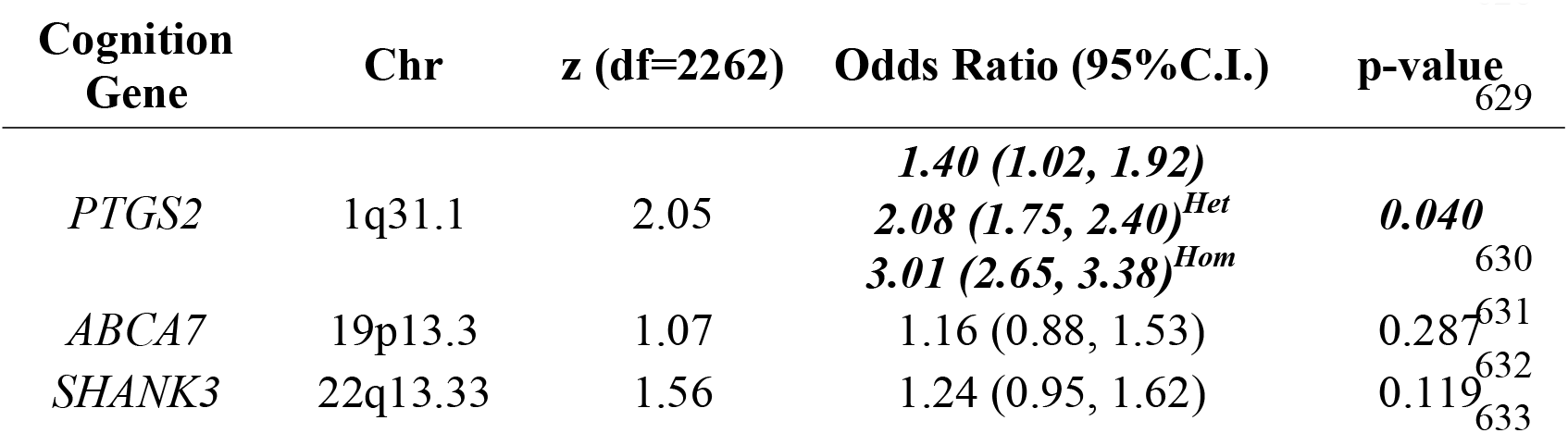
Association of cognition gene variants with intellectual disability in ASD.

**Table 2B.**
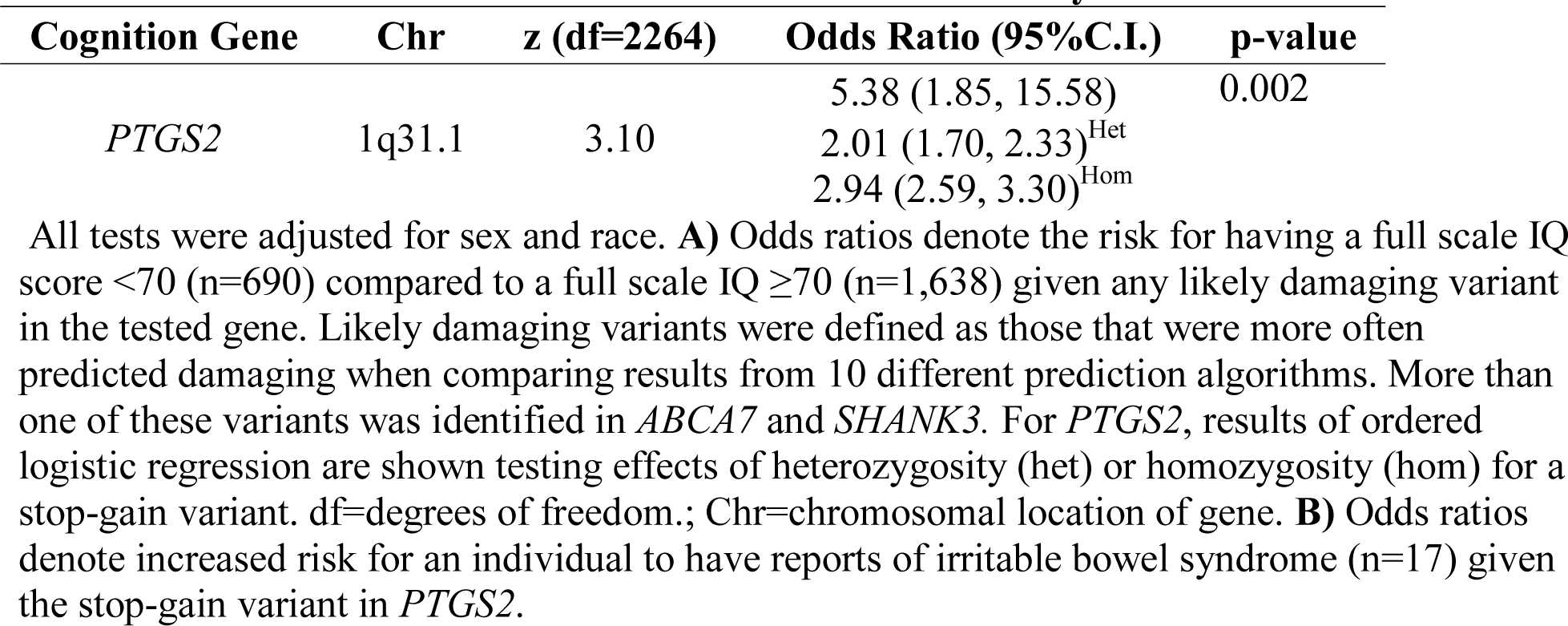
Association of *PTGS2* variant with irritable bowel syndrome in ASD.

## Discussion

### Novel Approach Identified Clinically-Relevant Genes to Prioritize for Functional Follow-up

Beginning with all 989 ASD candidate genes included in the December 2017 update of the AutDB Autism Informatics Portal, our approach identified a subset of 61 genes involved in cognition that were useful to defining a cluster of individuals with more severe teacher reported social impairment, lower IQ scores, and reduced daily living skills. We then identified three genes (i.e., *PTGS2*, *ABCA7*, and *SHANK3*) with likely damaging variants that were strongly associated with this ASD subgroup. This helped us to pinpoint the specific gene and variant that was associated with expression of important comorbidities in ASD, including intellectual disability and irritable bowel syndrome. In particular, a stop-gain variant in the *PTGS2* gene (rs200314986) encoding Cycloxygenase-2 (COX2) – a target for non-steroidal anti-inflammatory drugs (NSAIDs) – was more frequent in individuals with ASD compared to unaffected siblings. The support for *PTGS2* as a candidate gene for ASD resides in the results of a small gene-centric association study(*56*). As such, it is considered to have weak evidence for an association with ASD based on the cumulative strength of evidence for individual variants in that gene as defined in the AutDB Autism Informatics Portal. Our work provides additional support not only for a relationship between the *PTGS2* gene and ASD risk but also for increased risk of intellectual disability in ASD. Notably, the encoded enzyme is involved in serotonergic synaptic transmission and oxytocin signaling, which are known to be impaired in some individuals with ASD(*57-60*). While not this specific variant, there are 13 other pathogenic variants reported in this gene (https://www.ncbi.nlm.nih.gov/clinvar/) relating to developmental abnormalities. *PTGS2* is also considered a very important pharmacogene by PharmGKB and has strong implications for functional follow-up studies and eventual translation to improve clinical care(*61, 62*). There are a number of variants reported in this gene that have been shown to influence individual response to NSAIDs in the typically developing population(*61*). Given the evidence that long-term use of NSAIDs has been linked to gastrointestinal issues(*63*), we also tested for and observed that individuals with the *PTGS2* stop-gain variant had increased risk for reports of irritable bowel syndrome. It is possible that drugs that selectively inhibit COX-2, as well as traditional NSAIDs that target COX-2 and COX-1 (e.g., ibuprofen) may be less effective in individuals with this loss-of-function variant. This indicates that it may be useful to test for the rs200314986 variant in individuals with ASD to help improve treatment for pain and avoid exacerbation of gastrointestinal issues.

Variants in *ABCA7* were not strongly associated with ASD. Notably, there were 17 different likely damaging variants identified in this gene. This suggests that *ABCA7* may be more tolerant to loss of function mutations. We looked at loss intolerance scores (pLI), available via DECIPHER (https://decipher.sanger.ac.uk/) which assess the probability that a gene is intolerant to a loss of function mutation(*64*). These scores indicate that *ABCA7* may tolerate deleterious variants (pLI = 0.0). In comparison, there was only one stop-gain variant in *PTGS2* and three different variants (one splice-site and two CNVs) in *SHANK3*. *PTGS2* and *SHANK3* are predicted to be extremely intolerant (pLI for both genes = 1.00) to loss of function mutations.

Unexpectedly, *SHANK3* variants were associated with decreased risk for ASD. *SHANK3* is considered a strong candidate gene for ASD and haploinsufficiency of *SHANK3* is implicated in Phelan-McDermid syndrome which is often comorbid with ASD and characterized by delayed speech and intellectual disability(*65*). Notably, as we conducted gene-based tests our results were likely driven by a splice-site variant (rs150909992) that was observed to be heterozygous in 294 individuals, and not by the two CNVs. The splice-site variant was identified based on the VEP consequence from a previous assembly of the reference human genome (GRCh37.p13). This variant was not predicted by any of the other algorithms tested. In the most recent update of Ensembl (GRCh38.p10), this variant is no longer predicted to be a splice-site variant for a protein coding transcript of *SHANK3*. The transcript it affects corresponds to *SHANK3-202* which is now evidenced to encode a non-coding RNA. It is possible that this variant has no effect on the SHANK3 protein, which may explain why we did not see significant effects of having a likely damaging variant in *SHANK3* on risk for ASD or intellectual disability. It may instead have regulatory effects on other genes, as there is evidence that some non-coding RNAs are functional(*66*), and it is located in a promoter flanking region which is active in neuronal progenitor cells (ENSR00000147759). This is an excellent example of why it is important to consider that solely using the genetic location of the variant is potentially misleading in the ever-changing landscape of human genetics.

### Multiple Prediction Algorithms are Necessary for Efficient Identification of Damaging Variants

By evaluating damaging variant predictions from multiple algorithms, we were able to identify the variants (whether *de novo* or inherited, rare or common) in ASD risk genes with more evidence to be damaging to the encoded protein function. It is unclear what the optimal approach is for *in silico* prediction of the likelihood a genetic variant is damaging to the encoded protein product(*21, 33, 67*). Predictions from available tools vary widely when applied to the same variant as they employ different algorithms and use different training data to determine the accuracy of predictions(*68*). As such, it is highly advisable to combine predictions from multiple tools to assess the overall likelihood a variant is damaging(*69*). We observed that ∼13% of the SNVs and In/Dels that were expected to have a negative consequence on the encoded protein based on genetic location (i.e., the VEP prediction) were more often predicted to be benign by algorithms that incorporated additional information (e.g., the frequency of the variant in populations with no evidence of disease, the level of conservation of the genetic region across species). As such, if we had chosen to focus solely on VEP consequence predictions, we would have overestimated the likelihood of genetic dysfunction in the evaluated biological process. In addition, over half of the variants (52%) that were located in a genetic region that was likely to be damaging were not given predictions by any other algorithm. This is possibly because the variant being evaluated has not been observed in the populations that are used for training prediction algorithms. As such, it is currently difficult to determine the likelihood that an extremely rare variant is damaging without conducting functional follow-up studies. Only 0.14% of the variants that VEP predicted to be highly likely of damaging the protein product based on the location in the coding region of the gene were predicted to be damaging by all of the other nine prediction algorithms evaluated. Fortunately, as the field of *in silico* variant prediction continues to develop novel methods, focused on advances like mapping variants to three-dimensional protein structures(*70*), predictions should become more accurate and variant prioritization more efficient.

### Evidence of Intra-Individual Genetic Dysfunction in Multiple Biological Processes

The majority of the evaluated individuals had a variant in an ASD candidate gene that was predicted more often to be damaging compared to benign. By using these variants to calculate dysfunction in overall biological processes, we also observed that the majority of individuals had evidence of dysfunction in more than one process important to ASD etiology. The unique terms that were selected reflect validations of results from previous studies implicating genes involved in neural development, synaptic signaling, and chromosome packaging(*10, 11*). In addition, ASD risk genes were overrepresented in processes that encode the mental activities related to thinking, learning and memory (i.e., cognition) and regulate the difference in electric potential between the intra-and extra-cellular environments (i.e., regulation of membrane potential). While all of these processes had some degree of overlap in genes with likely damaging variants, there were also genes with variants that were uniquely assigned to only one process suggesting some genetic factors influencing these processes are distinct. Not surprisingly, individuals with more evidence for genetic dysfunction in development of the nervous system also had more evidence for dysfunction in the regulation of membrane potential and synaptic signaling. Dysfunction in genes influencing chromosome organization appeared independent from other processes. This provides additional support that mechanisms of chromosome organization may contribute independently from genes influencing neurological function to increase risk for ASD(*10, 11*). Notably, predicted dysfunction in cognition genes robustly identified a genetically-distinct subgroup of individuals with ASD. Many of these genes are evidenced to be involved in human cognition because they are implicated in intellectual disability, dementia, executive function, long-term memory, and a number of aspects of learning (for details see http://amigo.geneontology.org/amigo/term/GO:0050890). These genes may be particularly relevant to developing more comprehensive genetic screening panels for ASD. *Individuals with More Cognition Gene Dysfunction Have More Severe ASD Symptoms* The genetic subgroup defined primarily based on evidence of cognition gene dysfunction had increased severity of social impairment as measures via teacher reports on the Social Responsiveness Scale (SRS-TR)(*71*). Previous studies of families with more than one child diagnosed with ASD (i.e., multiplex) have observed that SRS scores are heritable, and linked to loci on a number of different chromosomes(*72-74*). SRS scores are observed to have differential distributions when comparing male and female individuals with ASD(*72*), and simplex versus multiplex families(*75*). Our results indicate that genetic factors influence social impairment measured via the SRS in simplex families, primarily in males. It is not clear why the observations are limited to teacher reports and do not extend to parent reports on the SRS-PR. We observed weaker correlations between parent and teacher reports on the SRS than has been previously reported(*76*). It is possible that these results reflect the highly variable symptom severity of the subjects in the SSC as concordance between teacher and parent reports is influenced by severity of ASD with higher concordance as ASD severity increases(*77*). Moreover, many studies have observed that parents rate their children as being more impaired compared to teachers possibly due to the context of the social setting in which the child is being observed(*78*). Notably, teacher reports on the SRS-TR were more correlated with social affect measured on the Autism Diagnostic Observation Schedule (ρ = 0.39) when compared to SRS-PR parent reports (ρ = 0.21) suggesting better agreement between teacher-reported and clinician-observed social impairment on average. Notably, PCA indicated that clinician-observed social/communication deficits measured via the Autism Diagnostic Observation Schedule (ADOS) were the largest contributors to the overall variability in quantitative ASD-related symptoms measured in the evaluated dataset and SRS-TR scores were among the top five.

Verbal and nonverbal IQ scores were also reduced in individuals with evidence of cognition gene dysfunction compared to those without. This was independent of social impairment measured via the SRS-TR, suggesting that individuals with more social impairment, and lower nonverbal and verbal IQ [as opposed to an ‘IQ split’(*79*)] have genetic differences compared to individuals with less social and intellectual impairment or discordance between these two measures (e.g., higher IQs with more social impairment). Previous studies have also observed that social deficits ascertained by the SRS are generally unrelated to IQ(*71, 80*).

### Limitations

Notably, many ASD candidate genes with likely damaging variants were assigned to more than one unique biological process. Therefore, to calculate scores for dysfunction in overall processes, genes with likely damaging variants were weighted to account for 1) the level of evidence supporting assignment of the gene to the biological process of interest, and 2) the proportion of the child terms of the unique biological process to which the gene was also assigned. For each gene assigned to a particular biological process, GO provides evidence codes that indicate the type of evidence supporting this assignment (http://www.geneontology.org/page/guide-go-evidence-codes). It is unclear what should be considered the most reliable sources of evidence supporting assignment of genes to GO Terms. While experimental evidence would be preferred, it is potentially biased, as this code will likely be assigned more often to genes that are directly evaluated for a role in the process of interest and not genes that have yet to be experimentally assessed for a role in the process. The majority of genes are assigned to terms based on computational predictions that have been observed to be generally reliable in the absence of experimental data(*81*). To avoid bias in gene process assignment, weights were calculated for each gene to account for the level of evidence it was correctly assigned to the process. It is possible that this approach under-or over-estimated the level of biological process dysfunction.

It is also possible that by focusing on currently implicated ASD risk genes we did not take into account all evidence for genetic dysfunction in a process. Future work aimed at understanding genetic contributions to overall process dysfunction, regardless of the underlying evidence of genetic risk for ASD may help detect more robust differences in ASD-related symptoms. In lieu of these potential limitations, the approach we developed helped identify the variants in ASD risk genes with more evidence to be damaging to the encoded protein function. This approach was also able to identify subsets of candidate genes with common underlying biology that are dysfunctional in individuals with ASD and related to differences in symptomatology. Notably, an inherited stop-gain variant in *PTGS2* was prioritized which has strong implications for functional follow-up studies and may be a novel treatment target. This work constitutes a translational bioinformatics approach beneficial to gleaning clinically-useful information from whole-exome data and could be adapted and applied to identification of clinically-relevant genetic factors for a number of complex human disorders.

## Supplemental Data description

Supplemental Data include six figures and eight tables.

## Acknowledgments

This work was supported by a National Library of Medicine grant K01-LM012870 [OJV] and the Simons Foundation (SFARI) [JSS, ZEW]. We are grateful to all of the families at the participating Simons Simplex Collection (SSC) sites, as well as the principal investigators (A. Beaudet, R. Bernier, J. Constantino, E. Cook, E. Fombonne, D. Geschwind, R. Goin-Kochel, E. Hanson, D. Grice, A. Klin, D. Ledbetter, C. Lord, C. Martin, D. Martin, R. Maxim, J. Miles, O. Ousley, K. Pelphrey, B. Peterson, J. Piggot, C. Saulnier, M. State, W. Stone, J. Sutcliffe, C. Walsh, Z. Warren, E. Wijsman). We appreciate obtaining access to phenotypic and genetic data on SFARI Base. Approved researchers can obtain the SSC population dataset described in this study (https://www.sfari.org/resource/simons-simplex-collection/) by applying at https://base.sfari.org.

